# speedingCARs: accelerating the engineering of CAR T cells by signaling domain shuffling and single-cell sequencing

**DOI:** 10.1101/2021.08.23.457342

**Authors:** Raphaël B. Di Roberto, Rocío Castellanos-Rueda, Fabrice S. Schlatter, Darya Palianina, Oanh T. P. Nguyen, Edo Kapetanovic, Andreas Hierlemann, Nina Khanna, Sai T. Reddy

## Abstract

Chimeric antigen receptors (CARs) consist of an extracellular antigen-binding region fused to intracellular signaling domains, thus enabling customized T cell responses against target cells. Due to the low-throughput process of systematically designing and functionally testing CARs, only a small set of immune signaling domains have been thoroughly explored, despite their major role in T cell activation, effector function and persistence. Here, we present speedingCARs, an integrated method for engineering CAR T cells by signaling domain shuffling and functional screening by single-cell sequencing. Leveraging the inherent modularity of natural signaling domains, we generated a diverse library of 180 unique CAR variants, which were genomically integrated into primary human T cells by CRISPR-Cas9. Functional and pooled screening of the CAR T cell library was performed by co-culture with tumor cells, followed by single-cell RNA sequencing (scRNA-seq) and single-cell CAR sequencing (scCAR-seq), thus enabling high-throughput profiling of multi-dimensional cellular responses. This led to the discovery of several CAR variants that retained the ability to kill tumor cells, while also displaying diverse transcriptional signatures and T cell phenotypes. In summary, speedingCARs substantially expands and characterizes the signaling domain combinations suited for CAR design and supports the engineering of next-generation T cell therapies.

## INTRODUCTION

Cellular immunotherapies against cancer have made substantial progress in recent years, with five FDA-approved chimeric antigen receptor (CAR) T cell treatments against hematological malignancies. These treatments rely on synthetic protein receptors that have been engineered for precise molecular recognition of a cell surface antigen (e.g., CD19 on the surface of B cell lymphomas). The infusion of autologous CAR T cells results in a cytotoxic response against tumor cells and, as in a classical immune reaction, this treatment can potentially result in persistent immunity, with CAR T cells recently observed in patients several years post-treatment [1]. However, success in this field has been difficult to replicate outside of hematological B cell malignancies. For example, cancers characterized by solid tumors, such as breast or lung, are more resistant to CAR T cell-mediated killing and have struggled to make progress clinically. Instances of remission, sometimes through antigen escape [2] have also highlighted the limitations of CAR persistence and of the monoclonality of the infusion product. Furthermore, strong adverse events, such as cytokine release syndrome (CRS) and transient neurotoxicity [3] are frequently associated with treatment and thus represent substantial safety concerns. Together, these pitfalls of CAR T cell therapies form substantial obstacles to their broader use against a wider range of cancer types.

In order to enhance CAR T cell responses against tumors, recent work has leveraged immunological mechanisms to counter the immunosuppressive microenvironment. For example, co-administration with checkpoint blockade inhibitors (e.g., monoclonal antibodies targeting PD-1/PD-L1) can activate tumor-infiltrating T cells [4]; or alternatively, “armoring” CAR T cells with additional genetic modifications to make them overexpress certain cytokines (e.g., IL-12, IL-23, etc.) can potentially sensitize tumors to cell-mediated cytotoxicity [5]. Furthermore, synthetic biology approaches that incorporate controllable domains [6-8] are being utilized to design CARs responsive to drugs or antigenic cues in order to tune the strength, duration and specificity of the inflammatory response. While promising, all of these strategies require additional drug compounds or genetic modifications, introducing further complexity in a therapeutic regimen that is already laborious and sensitive.

Recent advances in molecular immune profiling, such as single-cell sequencing and transcriptome analysis, are contributing important quantitative insights on CAR T cell-mediated responses and patient outcomes [9]. By going beyond methods for standard cell phenotyping (e.g., detection of surface markers by flow cytometry), transcriptional phenotyping can uncover gene expression patterns and metabolic pathways, or when used to track therapy progression and outcomes, they can help to identify predictors of therapeutic efficacy. A key observation from such studies is that the molecular components and design of a CAR has a major influence on the features of the transcriptional response [10]. A conventional CAR is rationally designed based on the fusion of modular gene elements: an extracellular antigen-recognition domain (frequently an antibody single chain variable fragment (scFv)), structural elements (a hinge, a transmembrane helix and peptide linkers), as well as one or more intracellular signal-transducing domains. The signaling domains are essential for linking the antigen-cell binding event to a cascade of intracellular molecular events, culminating in the expression of pro-inflammatory and cytotoxic genes. As would be expected, the signaling domains used can impact the type of response induced. However, almost all CARs in clinical trials to date combine the signaling domain CD3ζ of the T cell receptor (TCR) complex and the signaling domains of the co-receptors CD28 and 4-1BB [11]. Only a small panel of alternative signaling domains has been individually investigated clinically, and even fewer in parallel [10, 12, 13]. This lack of diversity is partly due to the low-throughput aspect of rational CAR design as well as the laboriousness of performing *in vitro* functional assays. Recently, two studies from Kyung *et al*. and Goodman *et al*. developed approaches to generate pooled CAR signaling domain libraries, which were screened by cell phenotyping (i.e., flow cytometry) to identify novel functional variants. [14, 15].

Here, we describe speedingCARs: single-cell sequencing of pooled engineered signaling domain libraries of CARs (Fig. 1). By exploiting the untapped diversity of natural immune signaling domains, we designed a modular signaling domain shuffling approach to generate a highly diverse library of 180 unique CAR variants, while maintaining a conserved scFv antigen-binding domain. The resulting CAR signaling domain library was integrated into primary human T cells by CRISPR-Cas9 genome editing. Pooled functional screening was performed by co-culturing with cognate antigen-expressing tumor cells, followed by single-cell RNA sequencing (scRNA-seq) and single-cell CAR sequencing (scCAR-seq). The combination of scRNA-seq and scCAR-seq enabled the high-throughput profiling of the multi-dimensional cellular responses, leading to the discovery of CAR variants displaying novel transcriptional signatures and T cell phenotypes. A subset of promising CAR variants were characterized with functional assays based on cytokine secretion, differentiation and cell-mediated cytotoxicity against tumor models, including 3D tumor microtissues. Thus, speedingCARs rapidly expands the diversity of the CAR synthetic protein family while simultaneously providing a deep transcriptional and phenotypic characterization of the newly-engineered variants.

**Figure 1:**
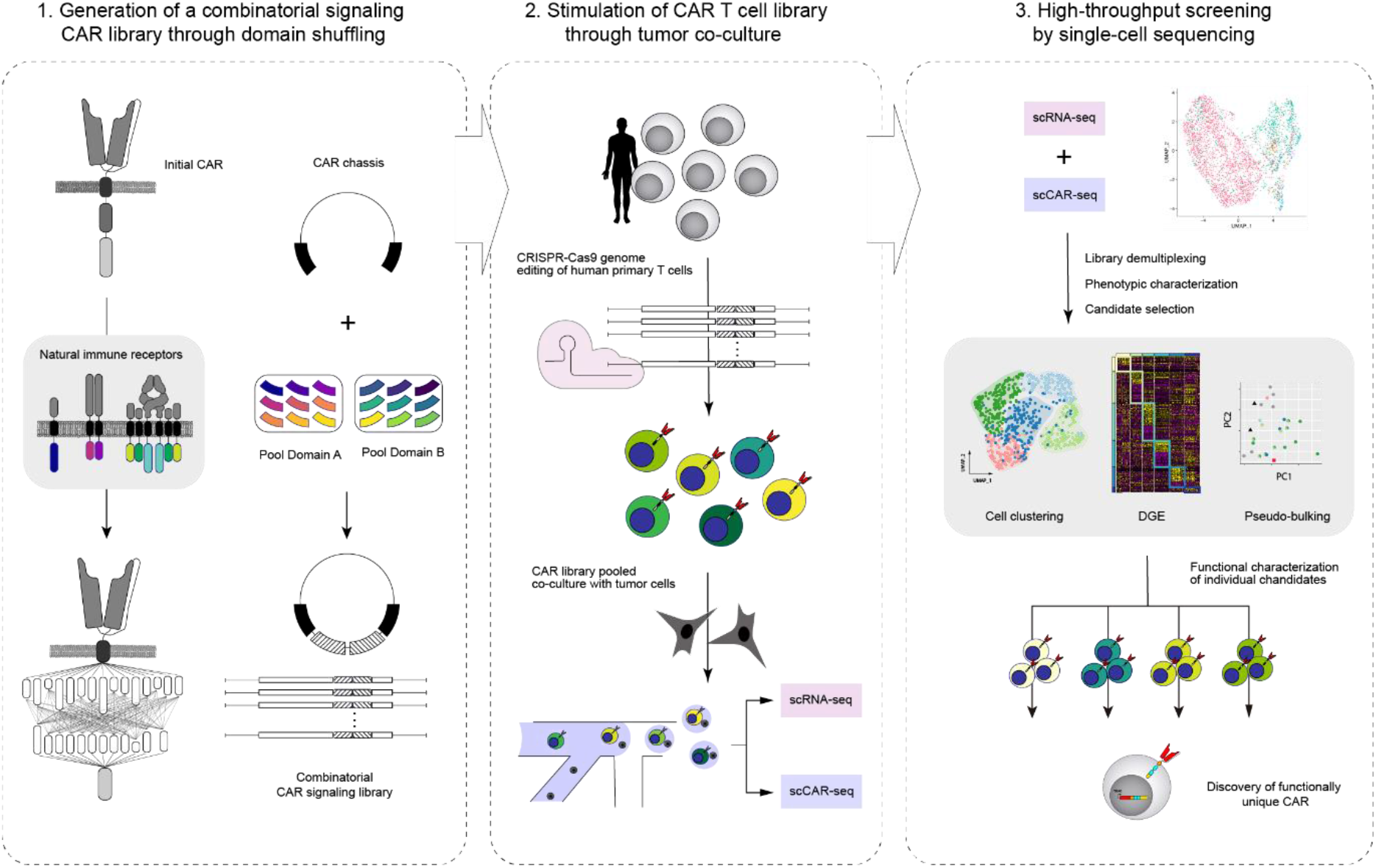
Schematic overview of speedingCARs: an integrated approach for the rapid engineering of CARs. **Left panel:** First, an initial CAR architecture is chosen based on functionality and encoded in a DNA plasmid. Next, one or more modular domains of the CAR are swapped for alternative natural immune intracellular signaling domain in a semi-random combinatorial fashion, resulting in a plasmid library of CAR variants with diverse combinations. **Middle panel:** The plasmid library of CAR variants is genomically integrated at the *TRAC* locus of primary human T cells via CRISPR/Cas9 genome editing. This ensures that each cell expresses a single CAR variant, and simultaneously deletes the endogenous TCR. A pooled library of CAR T cells is co-cultured in the presence of tumor cells expressing cognate antigen. **Right panel:** The functional screening of the pooled CAR T cell library is performed by scRNA-seq and scCAR-seq, revealing the transcriptional phenotype. This dataset is used to select promising variants for in-depth characterization with functional assays.

## RESULTS

### Design, generation and expression of a CAR signaling domain library in primary human T cells

In order to generate a library of CAR variants that could induce diverse T cell transcriptional phenotypes, we designed a modular cloning and assembly strategy for shuffling intracellular signaling elements (Fig. 2a). In a standard CAR, the intracellular region is composed of a CD3ζ signaling domain on the C-terminal end and co-receptor domains CD28 and/or 4-1BB on the N-terminal end (proximal to the transmembrane domain). CD3ζ is notable among other immune signaling proteins by the presence of three immunoreceptor tyrosine activation motifs (ITAMs), which play an important role in triggering downstream signaling events in response to receptor clustering. Although co-receptor domains such as CD28 and 4-1BB do not have ITAMs, their presence mimics the co-stimulation that accompanies TCR engagement with an activated antigen-presenting cell (APC), resulting in a more durable response.

**Figure 2:**
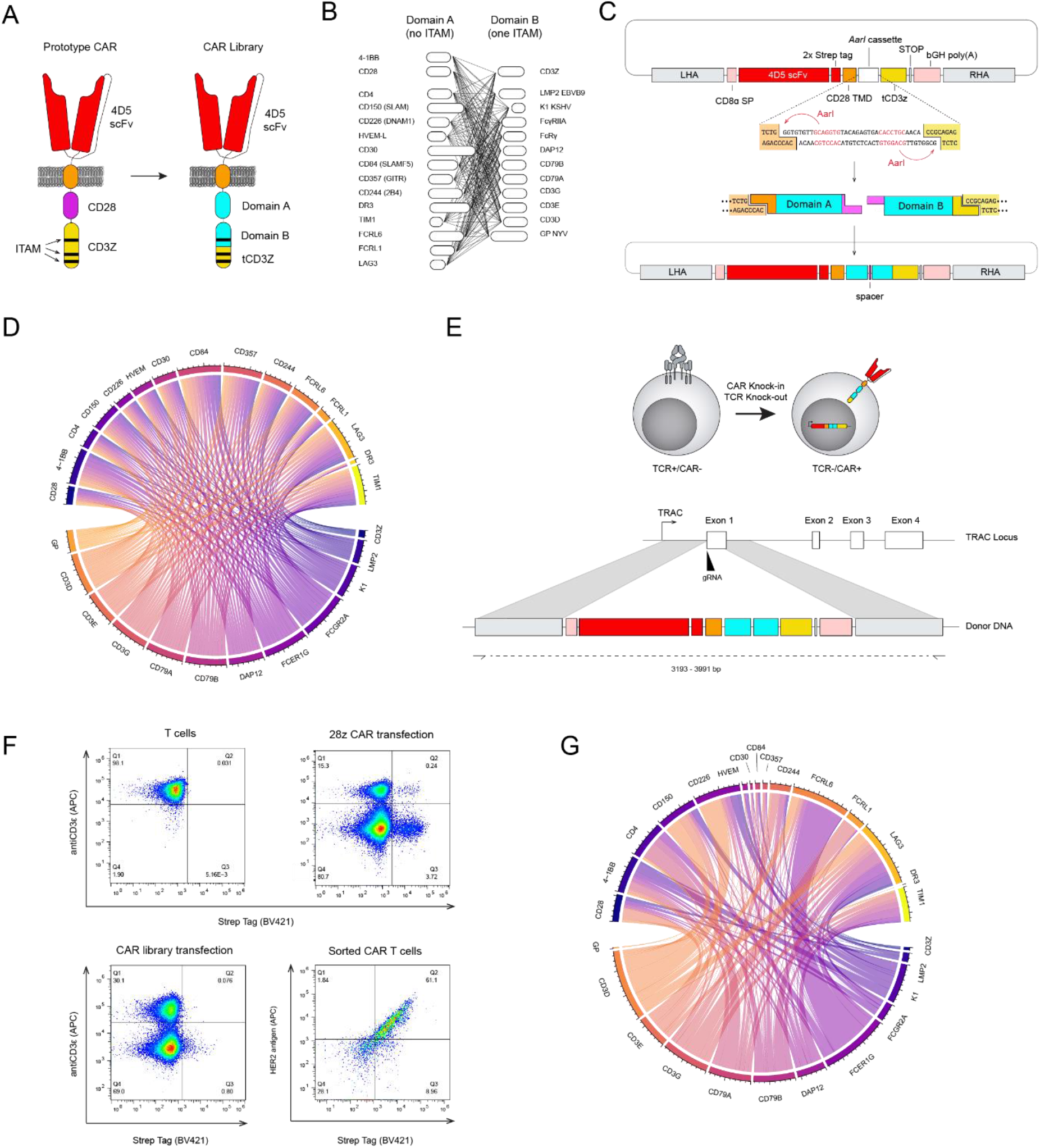
Combining modular domain shuffling and genome editing to express a library of CAR variants in primary human T cells. **A)** Schematic representation of CAR architecture for the shuffling of signaling domains. The CAR variant library is derived from an initial second-generation CAR featuring the intracellular signaling domains of CD28 and CD3ζ; the scFv (4D5) is based on the variable domains of the clinical antibody trastuzumab (specificity to the antigen HER2). The entire CD28 signaling domain and a segment of the CD3ζ domain are exchanged with signaling domains from two pools: Domain A and B, which possess either zero or one ITAM, respectively. A truncated CD3ζ (tCD3Z) possessing two ITAMs is retained. **B)** Combinatorial shuffling of Domain A and Domain B intracellular signaling domains yields a library of 180 possible CAR variants. **C)** Schematic representation of the cloning strategy for domain shuffling. A CAR chassis vector was designed encoding a conserved 4D5 scFv sequence, a cloning cassette and a part of CD3ζ. The cloning cassette replaced the co-signaling domain and the remainder of CD3ζ with outward-facing recognition sequences of the Type IIS restriction enzyme *AarI*. A restriction digest yields unique overhangs that are compatible with the ligation of domain from pool A and one domain from pool B, in that order (5’ to 3’). **D)** Long-read deep sequencing was performed following cloning and assembly of the signaling domain shuffling library. Circos plot shows that 179/180 possible combinations are present and in generally balanced proportions. **E)** Schematic of the strategy for targeted genomic integration of CAR library into the *TRAC* locus of primary human T cells by CRISPR-Cas9 HDR. **F)** Flow cytometry of human primary T cells before transfection, after transfection and after selection for CAR expression. **G)** Similar to panel C, circos plots display deep sequencing of the sorted library of CAR T cells, revealing a reduced diversity with unbalanced combinations of signaling domains.

Although recent work has highlighted some degree of plasticity in the number and configuration of ITAMs [16], our library design retained a total sum of three ITAMs to maximize functionality. To ensure this, we segmented the CD3ζ gene between the first and second ITAM and retained the two-ITAM segment for the CAR library. We then selected intracellular domains from a variety of immune co-receptors for inclusion into two pools of gene segments: domains A and B (Fig. 2b). The pool of domain A was generated from 15 receptors that have been described as providing co-stimulation in immune cell-cell interactions, ranging from the major contributors CD28 and 4-1BB to potentially more minor receptors, such as FCRL6 or CD150 and CD84 of the SLAM family. We also included lesser known or inhibitory receptors such as FCRL1, CD244 and LAG3 to investigate synergistic effects with co-simulators [17]. The pool of domain B consisted of 12 genes that each possessed a single ITAM. This included receptors such as FcγRIIA, DAP12 and CD79B which are expressed in diverse immune cells from both myeloid and lymphoid lineages, as well as T cell-specific genes involved in the TCR complex such as CD3G. In addition, we included three viral receptors, LMP2 (Epstein-Barr virus), K1 (Kaposi’s sarcoma-associated herpesvirus) and GP (New York virus), to investigate whether their non-human origin could lead to unique functional response profiles. Lastly, we also included the first segment of CD3ζ which would reconstitute the full intracellular domain upon assembly.

To shuffle and assemble the domain gene segments within the CAR genetic chassis, we used a Type IIS restriction cloning strategy [18] (Fig. 2c). In this method, customized restriction sites within a cloning cassette ensure that domains assemble in an orderly fashion: assembly of domain A on the N-terminal side and domain B on the C-terminal side results in the three ITAMs being proximal to each other. The two variable signaling domains are joined by a minimal linker. The remainder of the CAR chassis consisted of the following constant elements: (i) a secretion signal peptide from CD8α; (ii) an extracellular scFv based on the monoclonal antibody trastuzumab; (iii) a CD28-derived hinge and transmembrane domain; (iv) the partial segment of CD3ζ bearing two-ITAMs. Through the scFv, all CAR variants in the library have binding specificity for the oncogenic human epidermal growth factor receptor 2 (HER2), a clinically relevant target for CAR T cell therapy due to its prominence in many cancers [19]. Following cloning in a plasmid vector, the diversity of the resulting signaling domain library was assessed by deep sequencing, showing that the library possessed 179/180 possible combinations, with balanced representation (Fig. 2d).

To express the library of CAR variants in a physiologically relevant context, we performed genome editing with CRISPR-Cas9 on primary human T cells from healthy donors. Homology-directed repair (HDR) was used to target the CAR library for genomic integration at the TCR alpha chain (*TRAC*) locus (Fig. 2e). The precise integration of CARs at the *TRAC* locus has previously been shown to enhance tumor cell killing while conferring notable advantages over viral gene delivery (e.g., retrovirus, lentivirus): it ensures most T cells are monoclonal (only a single and unique CAR); it leads to transgene expression that is more consistent across cells and physiologically regulated; and it knocks out the endogenous TCR to avoid confounding effects [20]. Following genome editing, CAR T cells were isolated from non-edited cells by fluorescence-activated cell sorting (FACS) based on surface expression of a Strep tag and deletion of the TCR (Fig. 2f). The post-sort purity of the CAR T cell population and its ability to bind to soluble HER2 antigen was confirmed by flow cytometry, while genotyping validated targeted integration into the *TRAC* locus (Supplementary Fig. 1a and b). We also assessed the diversity of the library in this post-sort population by deep sequencing, which revealed reduced diversity (114/180 variants present) and unbalanced proportions of domain combinations (Fig. 2g). This is likely a reflection of the different stabilities and surface export efficiencies of each variant, and not a consequence of the HDR efficiencies owing to different sizes of the variants (Supplementary Figure 1c).

### Functional and pooled screening of CAR T cell library by single-cell sequencing

In order to achieve functional and pooled screening of the CAR T cell library, we performed co-cultures with tumor cells followed by scRNA-seq, which enabled us to profile transcriptional phenotypes and identify key gene expression signatures. This approach provides sufficient coverage of a CAR library of 180 unique variants by taking advantage of the current capacity of scRNA-seq (e.g., 10X Genomics, as used here) of 10^3^-10^4^ cells, which provides sufficient coverage of a CAR library of 180 unique variants. In addition, we designed scCAR-seq, a strategy to de-multiplex the CAR library from the resulting scRNA-seq data. For this, CAR single-cell barcoded transcripts found in the scRNA-seq cDNA product are selectively amplified and long-read amplicon sequencing (Pacbio) is used to capture the barcode identifiers that link the CAR signaling domain variant to a given cell (Supplementary Fig. 2).

In order to trigger activation of the CAR signaling domain variants, primary human T cells from two different healthy donors expressing the CAR library were co-cultured with the HER2-expressing breast cancer cell line SKBR3. To serve as a benchmark in subsequent analyses, we also co-cultured T cells bearing a standard CAR with the CD28-CD3ζ domain combination (abbreviated to 28z). Following co-culture (36 hours), CAR T cells were isolated by FACS and processed for scRNA-seq through the 10X Genomics pipeline. RNA transcripts were barcoded and amplified as cDNA to generate a gene expression (GEX) library suitable for deep sequencing. In parallel, scCAR-seq was performed and the resulting amplified transcripts were used for long-read sequencing.

### Identification of CAR-specific induced transcriptional phenotypes

Following pre-processing and quality filtering, scRNA-seq yielded a total of 6,421 CAR T cells. scCAR-seq allowed for CAR variant annotation of 70% of cells, from which 5% had to be discarded due to the integration of two different CAR constructs in the same cell. This resulted in a total of 4,199 correctly annotated cellular transcriptomes (averaging ∼3×10^7^ reads/cell) covering 98 unique variants from our CAR library (Fig. 3a). Despite a short co-culture time of 36 hours, which was intended to minimize more proliferative variants from becoming over-represented, evidence of clonal variant expansion was observed, particularly for the TIM1-CD79B variant. In order to do a clustering and gene expression analysis encompassing both over-represented variants (TIM1-CD79 and the 28z control) and less represented ones, we randomly subsampled up to 50 cells per variants. To confirm that this did not significantly affect clustering, 100 independent iterations of the subsampling and clustering algorithms were performed and the adjusted Rand index (ARI) was computed for each pair of partitions. This score, a measure of data clustering similarity, was consistently high, affirming that no specific subsample would likely change our results (Supplementary Fig. 3). The reduced dataset was used to do cell clustering and CAR variant enrichment analysis thereafter.

**Figure 3:**
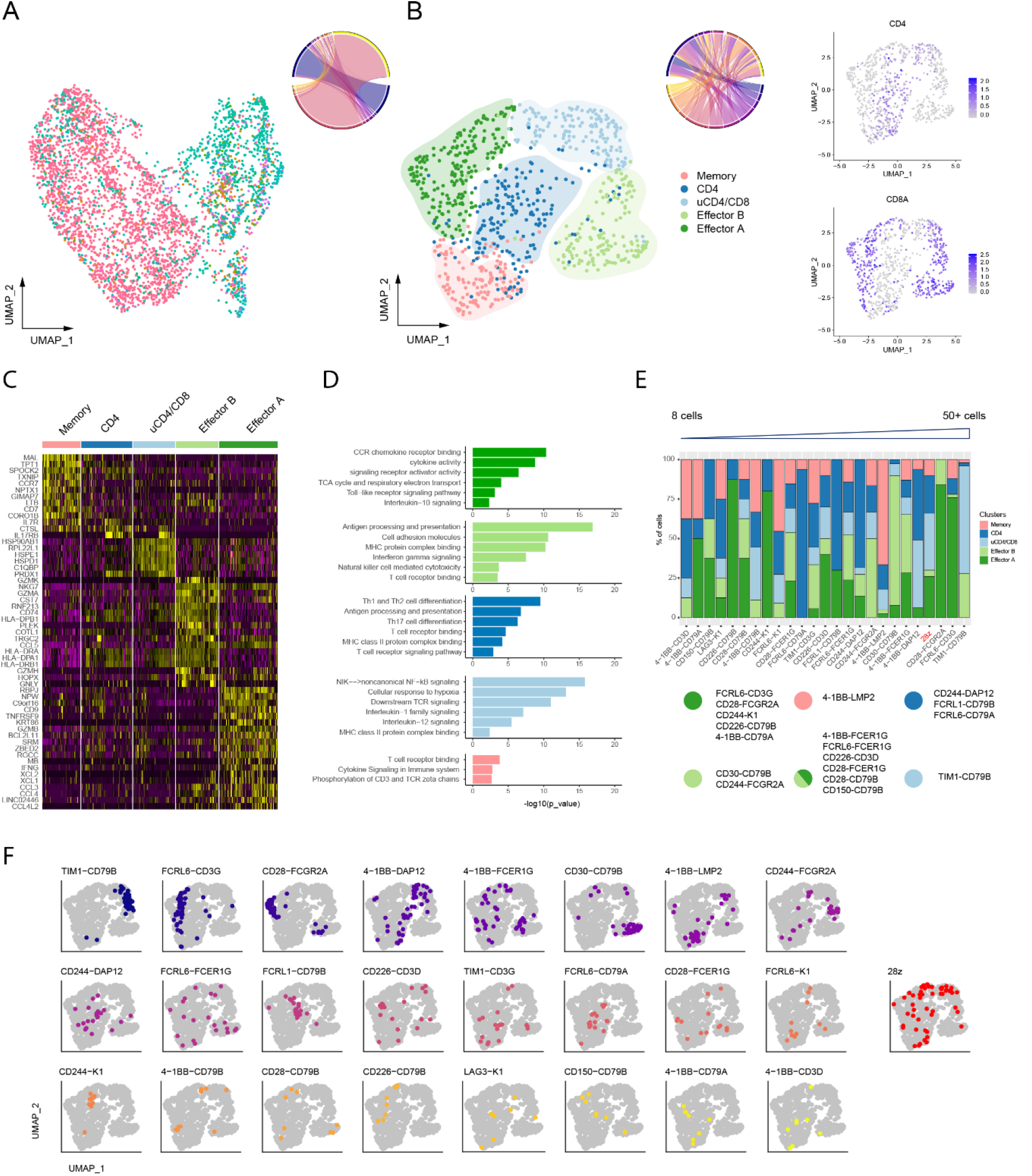
Unsupervised cell clustering classifies CAR variants based on distinct transcriptional phenotypes. **A)** UMAP embedding based on scRNA-seq of the pooled CAR T cell library following functional activation (co-culture with cognate antigen-expressing SKBR3 tumor cells. A total of 4,199 CAR T cells were successfully assigned to a unique CAR variant based on scCAR-seq. The dataset integrates cells from two independent donors and each colour indicates a different member of the CAR library. Circos plot illustrating the variant enrichment within the library. **B)** UMAP embedding and unsupervised cell clustering from scRNA-seq data of a subset of CAR T cells following functional activation. Shown are data from 823 CAR T cells resulting from randomly subsampling a maximum of 50 cells per variant (seed 42). The circos plot illustrates the variant enrichment within the library. On the right, UMAP feature plots show the distribution of CD4- and CD8-expressing cells. **C)** Heatmap of the differentially expressed genes between cell clusters in panel B. Only genes with at least 1.8 fold change are shown. **D)** Gene set enrichment analysis on differentially expressed genes between cell clusters in panel B. For each cluster, a selection of the most immunologically relevant gene sets is shown. **E)** Cluster enrichment observed for the top 24 most represented CAR variants. Variants are ordered by a confidence score, which is based on the number of available cells. CAR variants that have at least 50% enrichment on a given cluster are further assigned to six phenotypic groups. **F)** Distribution of sampled cells for each CAR variant within the UMAP embedding of panel B.

To determine whether CAR variants could trigger different functional T cell states upon encounter with tumor cells, we assessed the enrichment of each variant across T cell subsets. Unsupervised clustering identified five main clusters of T cell phenotypes, which were annotated based on CD4 and CD8 expression and differential gene expression of T cell marker genes (Fig. 3b and c, and Supplementary Fig. 4b): Memory, CD4, Unclassified CD4/CD8 (uCD4/CD8) and two CD8 effector clusters (Effector A and B). The Memory cluster was characterized by the enrichment of CCR7, IL7R, CD27, CD7, LEF1, SELL and KLF2. Effector CD8 cells were divided into Effector A and B clusters, both of which differentially expressed pro-inflammatory (CCL3, CCL4, CCL5 and CXCR3) and cytotoxic factors (NKG7, PRF1 and CST7) and some NK cell-associated receptors (KLRD1, KLRK1 and KLRG1). However, the Effector A cluster had higher levels of expression of canonical T cell effector genes (GZMB, IFNG, 4-1BB, RBPJ and ZBED2), while the Effector B cells differentially expressed other types of cytolytic factors (GNLY, GZMK and GZMA), cell migration related genes (LIME1 and CCR5) and >10 different HLA genes. CD4 and uCD4/CD8 clusters were less well defined by T cell marker genes. Some of the differentially expressed genes included CD40LG, IL17RB, IL13 and IL5. Cell cycle phase and cell donor was confirmed not to be a driver of cell clustering (Supplementary Fig. 4a). Clusters were further investigated using gene set enrichment analysis (Fig. 3c). For example, the Effector A cluster showed high cytokine and chemokine receptor binding activity and an active respiratory metabolism, the Effector B cluster displayed enrichment of T cell and NK cell activation features, cell adhesion and cytoskeleton rearrangement gene sets and the CD4 cluster was enriched on T cell activation and Th cell differentiation pathways. The uCD4/CD8 cluster possessed enrichment of genes predominantly not related to immune function; these were related to protein folding, cell cycle and cellular response to stress and hypoxia. The only immune-related gene sets showed non-canonical NF-kB and TCR signaling. After having defined the main cell clusters, CAR variant enrichment was used to further assess T cell function based on transcriptional profiles (Fig. 3e and f). Variants with at least 50% enrichment on a given cluster were assigned to a predictive phenotype.

### Interpretation of CAR variant-specific transcriptional signatures

Despite the general classification of cells into five clusters, there are still intra-cluster differences which do not capture the diversity of non-canonical T cell activation signatures that can be triggered by the new set of CAR cytoplasmic architectures. For this reason, we next examined the gene expression signature of each CAR variant individually using the full dataset of sequenced cells. A pseudo-bulking strategy was used to compare CAR variants with the standard 28z CAR (CD3ζ-CD28 signaling domains), including negative controls in the form of 28z CAR T cells without tumor co-culture (unstimulated) and unmodified T cells (not expressing CARs) co-cultured with tumor. Principal component analysis (PCA) revealed that CAR variants induced very diverse phenotypes and associated subgroups. Interestingly CAR variants assigned to an effector or a CD4 phenotype clustered closer to 28z CAR and away from the negative controls (unstimulated 28z and unmodified T cells) predicting a strong T cell activation response. On the other hand, CAR variants showing a high Memory and CD4/CD8 phenotype seemed to be transcriptionally closer to control samples, suggesting potentially a weaker T cell activation response (Fig. 4a). When examining the expression levels of the most variable genes, a similar clustering could be observed (Fig. 4b). Variants were divided in 4 different groups based on different levels of expression of pro-inflammatory (CCL3, CCL4, CCL5, CCL4L2, XCL1), cytolytic (NKG7, GZMB and GZMA) and additional regulatory factors (4-1BB, IFNG and IL13), allowing for inference of the functional properties triggered by each individual CAR variant. A different approach used to analyze individual CAR variants leveraged gene-set scoring. A score based on the expression of different gene sets was computed for each single cell allowing one to systematically screen CAR variants for specific T cell signatures (Supplementary Fig. 5). For instance, the FCRL6-CD3G CAR could be predicted to have improved cytotoxicity to 28z CAR, while having an increased memory phenotype and a higher tumor infiltrating potential. On the other hand, 4-1BB-LMP2 and LAG3-K1 CARs showed weak effector and cytotoxic potential.

**Figure 4:**
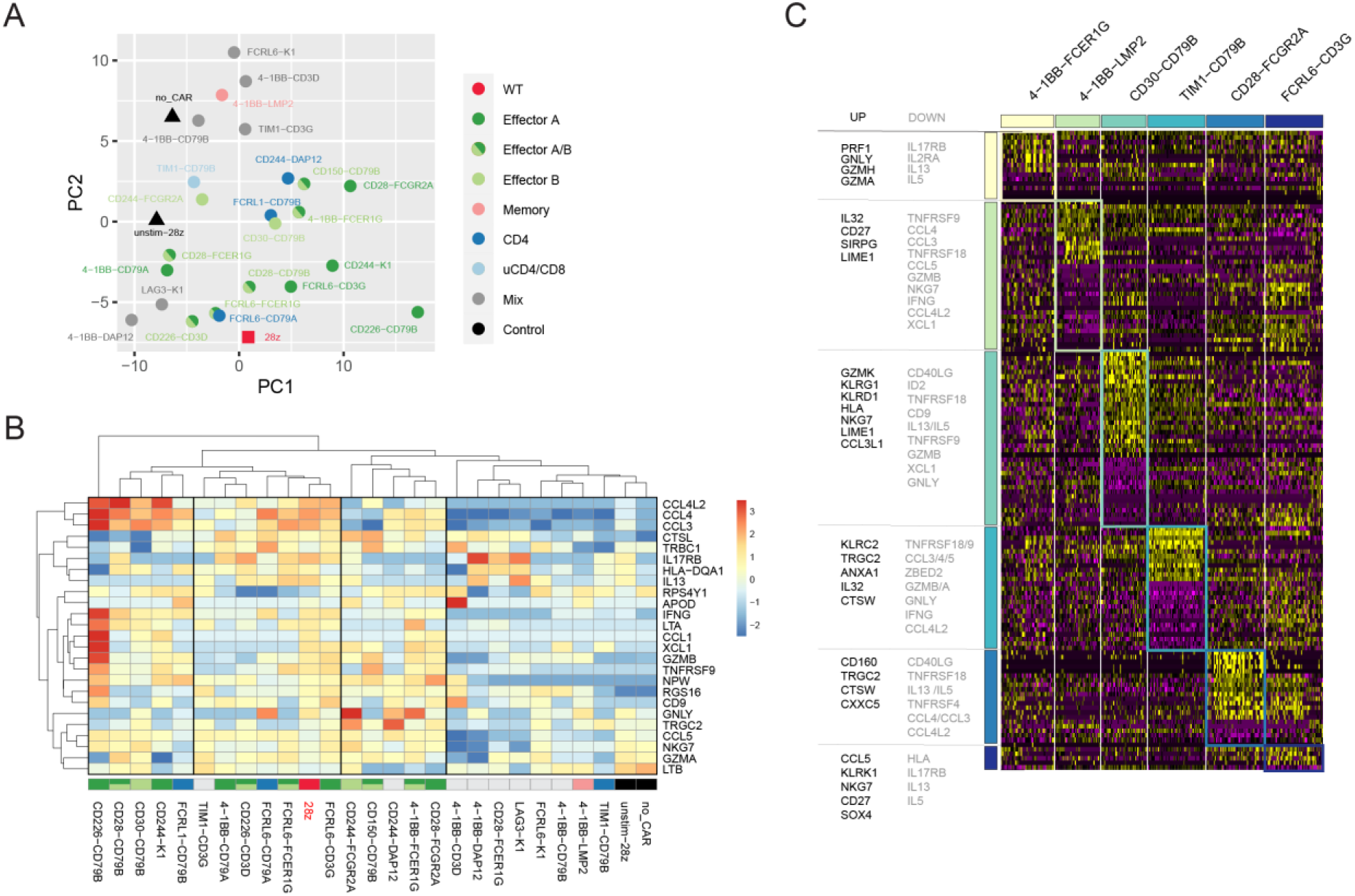
Specific gene transcriptional signatures are used to assess CAR T cell functionality. **A)** Principal component analysis (PCA) of pseudo-bulked scRNA-seq data from the 24 most represented variants of the CAR T cell library following functional activation. Also included are controls: 28z CAR T cells with and without tumor cell co-culture and non-genomically modified T cells with tumor cell co-culture. Samples are coloured following the annotation scheme described in Fig. 3e. **B)** Expression levels of the 24 most variable genes between pseudo-bulked scRNA-seq samples. Colour indicates normalised gene expression deviation from average. Genes and samples are ordered by hierarchical clustering and the sample annotation scheme described in Figure 3E is specified. **C)** Heatmap showing most differentially expressed genes between a selection of 6 different CAR variants and 28z CAR T cells following functional activation. Only genes with at least 1.7 fold change are shown both for over- and under-expression. The most immunologically relevant genes are highlighted for each CAR variant.

Following our survey of all CAR variants, we next selected a small panel of candidate variants from each of the cluster based annotation schemes and performed more detailed and direct comparisons with the clinically standard 28z CAR. Differential gene expression analysis on candidate CAR variants revealed phenotypes that may have promising clinical and therapeutic advantages (Fig. 4c). For example, the CAR variant 4-1BB-FCER1G showed enhanced secretion of perforin and cytolytic molecules, whereas variant CD30-CD79B exhibited a prevalent NK-like effector signature (NKG7, KLRG1 and KLRD1) and limited granzyme B secretion, providing a distinct but potent alternative to classic T cell mediated cytotoxicity. Also, the CD28-FCGR2A variant over-expressed CD160, which is known to play an important role in T cell-mediated cytotoxicity [21], whilst under-expressing CCL3 and CCL4, two factors known to promote CRS [22]. Notably, the FCRL6-CD3G variant displayed the closest transcriptional signature to 28z CAR, evidenced by the small number of differentially expressed genes, however it also presented a less terminally differentiated phenotype (higher expression of CD27) and increased expression of SOX4, a transcription factor associated with long term T cell persistence and suppression of Th2 differentiation [23, 24].

### In-depth functional characterization of selected CAR variants

Guided by the scRNA-seq data analyses, we selected five CAR variants and the clinical benchmarks 28z and 4-1BB-CD3ζ (BBz) for characterization with functional assays. The variants were selected on the basis of their notable transcriptional and signaling-associated phenotypes (Fig. 5a): variant A (CD28-FCGR2A) is associated with the Effector A cluster and showed reduced expression of CRS-associated factors; variant B (4-1BB-FCER1G) has high perforin expression and is present in both Effector A and B clusters; variant C (FCRL6-CD3G) displays an overall transcriptional phenotype similar to 28z but with potentially enhanced infiltration and persistence; variant D (CD30-CD79B) is associated with the Effector B cluster and has an NK-like signature; and variant E (TIM1-CD79B) has an apparently high proliferation rate. To perform these analyses on an individual basis, we generated CAR T cells for each variant by CAR genomic integration via Cas9 HDR in the *TRAC* locus of primary human T cells (as described above). As an additional control, we also sorted TCR knockout (TCR-KO) T cells [generated by Cas9-induced non-homologous end joining (NHEJ) in the *TRAC* locus]. Flow cytometry analysis of CAR expression showed that variants were generally expressed at slightly lower levels on the cell surface compared to the standard 28z CAR (Supplementary Fig. 6), which may be due to the use of the CD28 transmembrane domain. Next, following extended 5-day co-cultures with SKBR3 target cells, we profiled the secretion of an expansive panel of cytokines for each variant. The cytokines assayed were Th2-associated (IL-4 and IL-10), neutrophil-chemoattractant (IL-8) (Fig. 5b) or pro-inflammatory (IFNγ, TNFα, GMC-SF and IL6, Fig. 5c). Notably, the CAR variant A demonstrated a secretion profile very similar to that of 28z, which was characterized by the highest secretion of every cytokine tested with the exception of IL-10, where variant A was markedly lower. CAR variants B and C showed strong similarity with BBz, with all three occupying the middle range of cytokine secretion levels. CAR variants D and E formed a third similar group with lower cytokine secretion across the board, particularly for the Th2-associated IL-4 and IL-10. Lastly, all co-cultures with CAR T cells showed similar levels of IL-8, which is usually secreted by inflamed tissue. The presence of IL-8 in the T cell-free culture suggests that its origin was the SKBR3 cell line. Co-cultures with CAR T cells, and to a lesser extent with TCR-negative T cells, seems to have resulted in additional IL-8 release.

**Figure 5:**
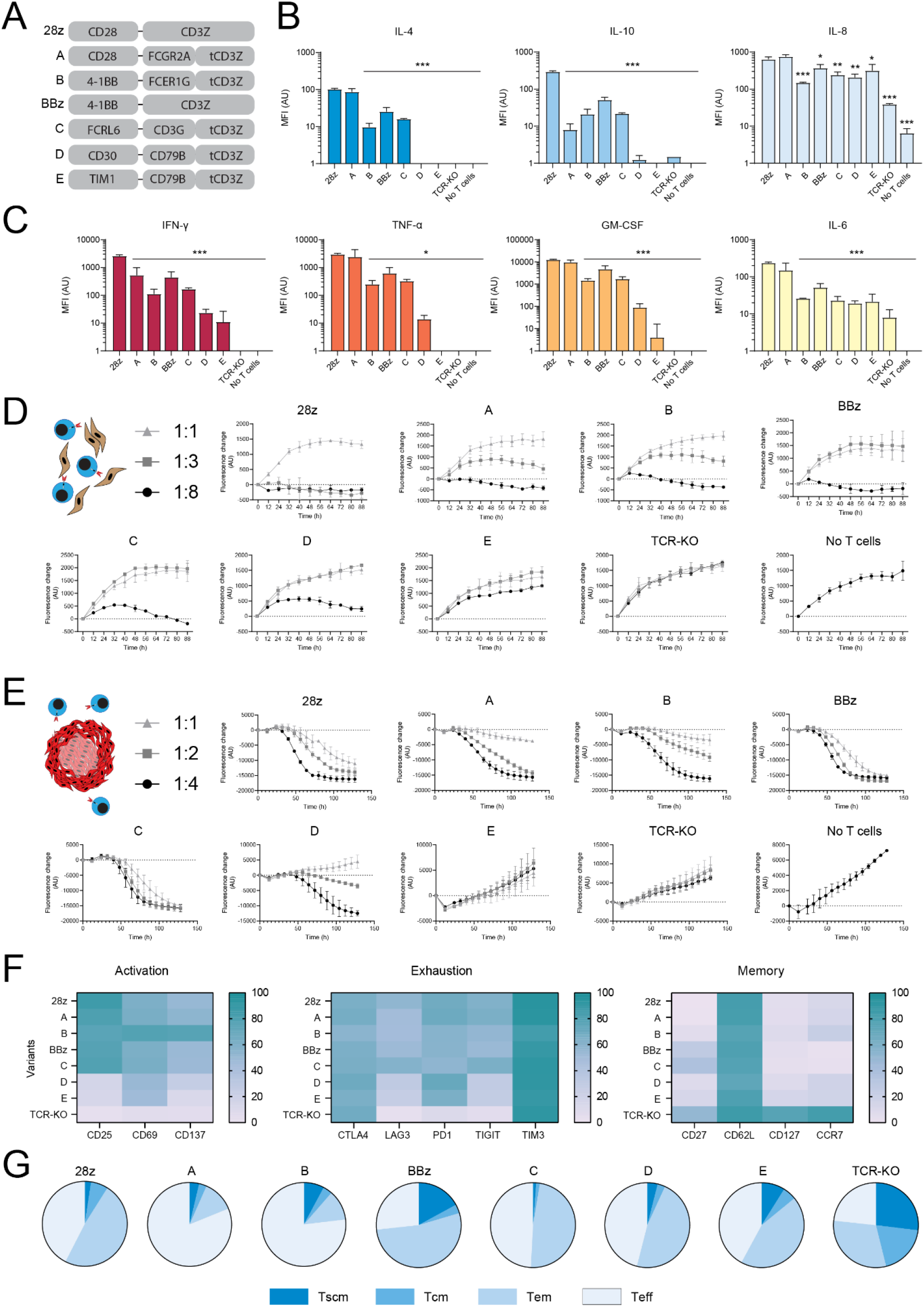
In-depth functional characterization of selected CAR variants confirms their diverse phenotypes. **A)** Five CAR signaling domain variants were selected for individual characterization along with the clinically used 28z and BBz CARs. **B) and C)** The relative levels of cytokines measured in the co-culture medium of CAR T cells and SKBR3 target cells suggest different immune activation profiles across variants. CAR T cells were co-cultured with SKBR3 cells in a 8:1 ratio and the medium was extracted by centrifugation for cytokine quantification by fluorescence-encoded multiplex bead assays. To assess significant differences between each variant and 28z, Dunnett’s multiple comparisons test was used with the following significance indicators: * P < 0.05, ** P < 0.001 and *** P < 0.0001. **D) and E)** The CAR T cell-mediated cytotoxicity of target cells proceeds at different rates depending on the signaling domain variants, the type of target and the CAR T cell:target cell ratio. CAR T cells were co-cultured with either SKBR3 adherent cells in a “sparse” culture in C) or with a single tumor spheroid of MCF-7 cells in D). Both target cell lines express GFP which was quantified over time by fluorescence microscopy to assess target cell death. The values represent the difference in GFP intensity from the first time point. **F)** Heatmap of the proportion of CAR T cells expressing surface markers of activation, exhaustion and memory for each variant. The CAR T cells co-cultured for the cytotoxicity experiment in C) were used in a multi-parameter flow cytometry assay. **G)** Pie charts of the T cell differentiation subsets for each variant following co-culture. Based on the surface expression of CD45RA, CD45RO and CD27. In all panels, TCR-KO refers to T cells without a TCR and error bars represent the S.E.M.

Next, we measured the cytotoxic capacity of the selected CAR variants for tumor cell control and elimination through live-cell imaging. As target cell lines, we used SKBR3 and MCF-7, the latter of which is another breast cancer cell line expressing HER2, albeit at lower levels [16]. To track the cytotoxic activity via cancer cell death, we used Cas9 HDR to integrate a GFP expression cassette at the *CCR5* genomic locus of SKBR3 and MCF-7 cells (Supplementary Fig. 7a). After three rounds of iterative sorting, the resulting populations showed stable expression of GFP in over 90% of cells (Supplementary Fig. 7b-e). For SKBR3, we tracked the total fluorescence of standard “sparse” co-cultures with the selected CAR T cell variants at various ratios (Figure 5d). At the lowest ratio of CAR T to tumor cells (1:1), no variant could control SKBR3 expansion. At the intermediate ratio (3:1), the standard 28z CAR along with variants A and B could control or eliminate SKBR3 expansion, however this was not the case for the other clinically established BBz CAR. The CAR variants C and D could only control target cell expansion at the highest ratio (8:1), while variant E only had a slight effect. As an alternative cytotoxicity model, we sought to challenge the CAR variants against three-dimensional cancer microtissues (spheroid structures). We established conditions under which the GFP+ MCF-7 cell line forms spheroids (Supplementary Fig. 7f and g), which were then cultured with CAR T cell variants at various ratios. Within this spheroid setting, we observed marked differences not only in outcome, but also in the rates of tumor cell spheroid killing (Fig. 5e). Variants A and B reduced spheroid mass at all ratios but substantially slower than the standard 28z and BBz CARs. Variant C performed similarly to the BBz CAR, showing rapid elimination at all ratios, making it much more effective in the 3D setup compared to the sparse culture model. In contrast, variant D displayed the slowest cytotoxicity, while variant E once again failed to control spheroid growth at all ratios.

In order to determine the extent to which CAR variants influence T cell differentiation programs, we performed multi-parameter flow cytometry on 16 surface markers, including those characteristic of memory, activated and exhausted T cell phenotypes (Fig. 5f and Supplementary Fig. 8a and b). Following 5-day co-cultures with SKBR3 cells, surface expression profiles revealed notable differences across variants despite broadly similar CD4/CD8 ratios. For activation, the expression patterns of CD25 and CD69 generally matched what had been observed in terms of cytotoxicity, with variant B standing out for its additional strong expression of CD137. For exhaustion, all variants showed relatively similar expression levels of CTLA4 and TIM3. Variants B and C showed slightly lower PD-1 expression compared to CD28z and variant A, while variant C showed elevated levels of LAG3 and TIGIT. Memory markers were generally similar with slightly higher CD27 levels in BBz and variant C. Some of these observations remained when gating on CD8+ cells only (Supplementary Fig. 8c). Lastly, we used CD45RA/CD45RO expression and CD27 to subset CAR T cells into the four standard T cell differentiation sub-populations: T stem cell memory (Tscm), T central memory (Tcm), T effector memory (Tem) and T effector (Teff) (Fig. 5g). The proportion of subpopulations for each of the variants were largely similar to one another with some notable exceptions: i) high Teff / Low Tscm in variant B; 2) high Tcm and low Tem in variants A and C; and 3) high Tem and low Tcm in variant D. It should be noted that CD27 is but one possible marker used to define these subsets; alternative ones include CD62L and CCR7 and a comparison between these definitions is provided (Supplementary Fig. 8e).

## DISCUSSION

Here, we developed speedingCARs, an integrated method that combines signaling domain shuffling and single-cell sequencing to expand the range and functional profiles of CAR T cells, which may eventually enable enhanced cell therapies. Our approach, which is designed for rapid and productive CAR engineering, utilizes two important concepts. First, CARs were conceived with modularity in mind, making them especially suitable for the combinatorial shuffling of signaling domains derived from a wide range of receptor types. Second, scRNA-seq is currently feasible at an appropriate scale for screening a pooled library of CAR variants in order to identify transcriptional phenotypes with unique T cell activation profiles. To maximize translatability, we introduced the signaling domain-shuffled library into primary human T cells and triggered CAR activation through a co-culture with tumor cells expressing cognate antigen (HER2). This unique strategy allowed us to identify functional CARs with previously unused intracellular signaling domain combinations. These candidates exhibit properties that are uncommon in current standard designs and, thus may lead to new applications for CAR T cells.

Among natural proteins, domain shuffling is thought to be a major evolutionary driver [25]. By definition, a domain’s structure is modular and its function is portable. DNA translocation events that carry a domain-coding sequence into another gene can thus be well-tolerated from a functional standpoint. A domain might synergize with its neighboring domains, e.g. a kinase domain joining a binding domain, potentially creating a new pathway branch [26, 27]. Signaling proteins are especially likely to be modular, leading to their embedding in complex networks of protein-protein interactions. With their signaling domains typically taken from T cell immune receptors, CARs are no exception. While the understanding of natural T cell signaling has benefited from decades of research, there is little knowledge on the effects of mixing and matching signaling domains in synthetic receptors such as CARs. Efforts to date have selected domain combinations largely by trial and error and from a small pool of known effectors [10, 12, 13]. Inspired by natural evolution, the speedingCARs method relies on random domain shuffling to generate a library of all possible combinations from which functional pairings can be identified.

While domain shuffling constitutes a powerful method for rapidly engineering new diversity in a protein, this can make the choice of screening strategy a challenging ordeal. Previous work for CAR engineering has relied on functional screening based on the expression of single reporter genes or proteins (e.g., IL-2, NFAT, CD69) in immortalized cell lines [15, 28-30]. While these approaches enable high-throughput screening of CAR libraries, they are limited by their uni-dimensionality: they generally reveal only a single aspect of the effector response and do not capture the full complexity of the deeply interconnected signaling network of T cell activation. Furthermore, immortalized cell lines do not fully recapitulate primary T cells, reducing the translatability of the screening results. Rather, to obtain multi-dimensional and translatable CAR functional profiles, primary T cells and RNA-seq offer a powerful alternative. The use of scRNA-seq in particular is fast becoming an important tool in the characterization of CAR T cells before and during their clinical evaluation [9], and it is especially suited for the screening of a pooled library of engineered cells. Recent work by Roth *et al*. demonstrated how scRNA-seq can be used to capture the transcriptome and the corresponding identity of a library member in engineered cells [31]. In their approach, a pooled library of 36 knock-in genes was screened in combination with a NY-ESO-1-specific TCR to find transcriptional phenotypes that could enhance the induced T cell response. As the capacity of scRNA-seq continues to grow, larger libraries can be screened in this way. Here, we harnessed current scRNA-seq capabilities to screen a library of 180 possible CAR signaling variants and directly identify unique transcriptional phenotypes of T cell activation. Our method circumvents the need for engineered cell lines and reporter genes, as it retains a robust throughput while greatly enhancing the translatability of the screening results.

By shuffling signaling domains from diverse origins, we aimed to identify CARs that promote unique transcriptional phenotypes, as well as domains that may be associated with novel immunological profiles and function. Indeed, we found significant phenotypic diversity in our CAR library, especially with genes related to effector and cytotoxic function, memory differentiation, Th1/Th2 classification and tumor infiltration. These unique profiles may prove valuable in specific contexts where maximal cytotoxicity is not the only sought-after property. For instance, in clinical settings, BBz CARs have sometimes proven to be superior to the 28z combination with respect to T cell exhaustion and persistence *in vivo* [32, 33], or showing a reduced incidence of adverse events [34]. The cytokine and cytotoxicity functional assays we performed suggest that all of our selected CAR variants have some ability to trigger a response compared to the controls, and some resemble BBz in their profile. Furthermore, some signaling domains appear more productive than others. For instance, in terms of representation in the transcriptomics screen, the prominence of combinations featuring CD79B may reflect an ability to induce proliferation. Following selection, we found that variants A, B and C, all featuring Fc or Fc-like receptor domains, induce the most pro-inflammatory cytokine secretion and can control or eliminate tumor cells. FcεRIγ, present in variant B, was one of the two original signaling domains used in early CARs, the other being CD3ζ [35]. Our results suggest that other Fc domains and combinations may be worth testing further. Likewise, CD30, a member of the tumor Necrosis Factor Receptor (TNFR) superfamily may be responsible for the enhanced response of variant D versus E. Little is known on the exact function of this receptor, though its interaction with TNFR-associated factors (TRAF) and subsequent signaling [36] may have played a role here. In their flow cytometry-based CAR screening method, Goodman *et al*. also identified TNFR superfamily members as conferring functional enhancements [14]. Lastly, we note that the presence of the CD28 and 4-1BB signaling domains among selected variants confirm their important co-stimulatory properties. However, other CAR constructs incorporating these domains (i.e. 4-1BB-LMP2 and 4-1BB-DAP12) showed poor T cell effector potential in the transcriptomics data, affirming that the contribution of all signaling domains in a CAR matters for a given T cell phenotype.

Many of the pitfalls of CAR T cells are being addressed with complementary solutions such as additional gene editing to enhance the T cell immune response or the co-administration of immunomodulating compounds [4-8]. These solutions risk adding layers of complexity to what remains a challenging treatment to administer. Ideally, a single genome-integrated CAR would suffice, as in currently approved regimens, but different tumors may require slightly different approaches, making it challenging to find an optimal construct for a given situation. The speedingCARs method offers a path to this next step of personalized precision medicine. Likewise, there is now growing interest in using CARs against other conditions, such as viral infections [37, 38] or autoimmune disease [39, 40]. Likewise, CAR-expressing natural killer (NK) cells and macrophages are also being investigated [37, 41, 42]. For subtype-specific CAR T cells (such as CAR Tregs) or innate cell effectors, it remains unclear whether CD28, 4-1BB and CD3ζ are truly optimal. While our study design was conservative in that the majority of the signaling domains we shuffled came from pro-inflammatory T cell-associated effectors, our approach could be expanded to include larger sets of domains or sets composed exclusively of specific protein classes (e.g. inhibitory, innate immunity-linked, etc.) This could yet expand the possible phenotypic space of CARs to previously unreached areas.

## METHODS

### Cloning of the library

The CAR signaling domain library was constructed in a DNA plasmid vector using a Type IIS restriction enzyme cloning strategy, as previously described [18]. Briefly, the vector was designed with a cloning cassette within a CAR chassis, as illustrated in Fig. 2b. In this chassis, the CD3ζ signaling domain was segmented, retaining amino acids 100 to 164 (amino acids 52 to 99 were used in the pool of domain B). The vector was digested with the Type IIS restriction enzyme AarI (Thermo Fisher) for 4 hours at 37°C and treated with Antarctic phosphatase (NEB) for 30 min at 37°C. The signaling domains were amplified from synthetic DNA gene templates (Twist Bioscience) with primer pairs F1/R1 or F2/R2 and digested with AarI. An equimolar mix of all domain fragments was prepared for ligation into the digested CAR chassis vector with T4 ligase (NEB). The ligated plasmids were transformed and amplified in chemically-competent *E. coli* DH5α cells (NEB).

### Primary human T cell isolation and culture

To obtain human primary T cells, buffy coats from healthy donors were used (Blutspendezentrum, University of Basel) and the peripheral blood mononuclear cells (PBMCs) were extracted from a Ficoll gradient. The T cells were isolated using the EasySep human T cell isolation kit (Stemcell) and activated with human T-activator CD3/CD28 Dynabeads (Gibco) at a bead:cell ratio of 1:1. Activated T cells were cultured in X-VIVO 15 (Lonza) supplemented with 5% fetal bovine serum (FBS), 50 µM β-mercaptoethanol, 100 µg/mL Normocin (Invivogen) and 200 U/mL IL-2 (Peprotech), thereafter referred to as T cell growth medium. After 3 days, the beads were magnetically removed.

### Primary human T cell genome editing

We adapted our previous CRISPR/Cas9 genome editing protocol [28] to introduce CAR genes at the *TRAC* genomic locus. Briefly, the ribonucleoprotein (RNP) particles were assembled by first duplexing the CRISPR RNA (crRNA) and trans-activating CRISPR RNA (tracrRNA) (IDT) through co-incubation at 95°C for 5 minutes and cooling to room temperature. The duplexed RNA molecules were then complexed with 25 µg (153 pmol) of Cas9 protein (IDT) at room temperature for 20 minutes. For HDR with double-stranded DNA repair template, the primers F3 and R3 were used to amplify the CAR gene and homology arms with truncated Cas9 target sequences (tCTS) as described in [43]. The product was purified and 2 µg were added to the RNP and diluted in 100 µL P3 nucleofection buffer (Lonza). This mixture was nucleofected with 2×10^6^ stimulated human primary T cells using the 4D-Nucleofector (Lonza) with the program EO-115. The cells were then immediately diluted in 600 µL T cell growth medium.

### Cancer cell line culture and genome editing

The cell lines SKBR3 and MCF-7 were cultured in Dulbecco’s Modified Eagle Medium (DMEM) (Gibco) supplemented with 10% FBS, 1% penicillin-streptomycin (Gibco) and 50 mg/mL Normocin (Invivogen), thereafter referred to as cell line growth medium. CRISPR/Cas9 genome editing in these cell lines was performed with RNP particles as described above with the following differences: the gRNA was specific for *CCR5* and taken from [44]; the nucleofection buffer was Dulbecco’s phosphate-buffered saline (DPBS) (Gibco); the nucleofector protocol was EO-117 for SKBR3 and EN-130 for MCF-7; and the cells were diluted in cell line growth medium.

### Library sequencing

We used next-generation sequencing to characterize the diversity of the CAR signaling domain library. To examine the library at a plasmid level, we amplified the shuffled signaling domains using a PCR reaction with primers annealing to flanking sequences (F4 and R4; Supplementary Table 2) and purified the resulting product ranging between 304 and 1096 bp. To assess the library diversity following genome editing, we performed a two-step PCR. First, genomic DNA was extracted from 10^4^ to 10^5^ CAR-expressing T cells using QuickExtract buffer (Lucigen). The resulting product was used as a DNA template for a first PCR amplification reaction using F5 and R5 primers (Supplementary Table 2) to produce a long amplicon which confirmed *TRAC* locus genomic integration. This product was then used as DNA template for a second PCR amplification using primers F4 and R4. The final amplimers were purified and sequenced externally by Illumina paired-end sequencing (GENEWIZ).

### FACS of CAR expression and binding

Analytical and sorting flow cytometry protocols to confirm genomic integration of CAR constructs were described before and adapted here to primary human T cells [5]. The knock-out of the T cell receptor (TCR) was assessed with the absence of signal after staining 1:200 with APC-labeled anti-CD3ε (Biolegend). Each CAR construct contained a Strep tag which allowed for a two-step staining to validate successful knock-in; a 1:200 biotinylated anti-Strep tag antibody (GenScript) treatment was followed by a 1:400 Streptavidin-BrilliantViolet 421 conjugate (Biolegend). Similarly, HER2 binding was confirmed with 2.5 µg/mL soluble HER2 antigen (Merck) and subsequent 1:200 APC-labeled anti-HER2 antibody (Biolegend) incubation. Cells were washed in cold DPBS and kept on ice until analysis. CAR T cells were sorted into room temperature T cell growth medium and maintained for 5 days to recover before co-culture assays.

### Single cell sequencing and data analysis

Previously sorted library CAR T cells (rested for 8 days following bead removal) were co-cultured for 36 hours with the high HER2-expressing tumor cell line SKBR3. An E:T ratio between 0.2-0.5 was used to maximize the contact of CAR T cells with their target antigen. Immediately after co-culture, CAR-T cells were sorted by FACS and single-cell RNA sequencing was performed using the 10X Genomics Chromium system (Chromium Single Cell 3’ Reagent Kit, v3 chemistry; PN-1000075) following manufacturer’s instructions. In short 4000-10000 cells were resuspended in PBS and loaded into a chromium microfluidics chip. Following GEM formation, reverse transcription and cDNA amplification, 25% of the sample was used for 3’ gene expression library preparation including the incorporation of Chromium i7 multiplex indices (PN-120262). The resulting transcriptome libraries were sequenced using the Illumina NovaSeq platform. scRNA-seq data was generated from two individual donors. scRNA-seq data was generated from two individual donors.

### Single cell CAR sequencing (scCAR-seq)

In order to de-multiplex the CAR T cell library within the scRNA-seq data, a scCAR-seq strategy was developed (adapted from [31]; Supplementary Figure 2). Using 40 ug of 10X cDNA, the cytoplasmic region of the 10X barcoded pooled barcoded CAR cDNA molecules was amplified using KAPA-HIFI, a Read1-p5 primer (F5) and a customized Strep-Tag specific Read2 primer (R5). Following a 10 cycle PCR (95C for 3’, [98C for 20”, 67C for 30”, 72C for 60”] ×10, 72C for 2’) and a X0.65 SPRI bead DNA clean up (AMPure XP, Beckman Coulter) a second PCR using a p5 primer (F6) and a i7-Read2 primer (Chromium i7 Multiplex Kit, 10X Genomics, PN-120262) was used to further amplify the genetic material for 15 cycles (95C for 3’, [98C for 20”, 54C for 30”, 72C for 60”] x15, 72C for 2’). CAR amplicons were then sequenced using PacBio SMRT sequencing platform and Biostrings R package was used to assign CAR variants to each 10X single-cell barcode.

### Analysis of scRNA-seq data

The raw scRNA-seq data was aligned to the human genome (GRCh38) using Cell Ranger (10x Genomics, version 3.1.0) and downstream analysis was carried out using the Seurat R package (version 4.0.1; [45]). Low quality cells were removed based on the detection of low number of UMIs (nFeature_RNA > 200), high number of RNA molecules (nCount_RNA < 150 000) and high percentage of mitochondrial genes (Percentage_MT < 20% of total reads). scCAR-seq results were then used to assign CAR variants to each sequenced cell, obtaining an assignment rate of 70% of cells, and only cells assigned to a single CAR variant (65%) were used for downstream analysis.

After QC and CAR T cell assignment a total number of 3227 stimulated library CAR T cells, 972 stimulated 28z CAR T cells, 5439 unstimulated 28z CAR T cells and 3027 TCR-KO T cells were obtained. Each dataset was log normalized with a scale factor of 10 000 and sample integration was performed applying the reciprocal PCA seurat pipeline using 2000 variable integration features. Integrated data was then scaled regressing out cell cycle genes and dimensionality reduction was done using the *RunPCA* function. *FindNeighbors* and *Findclusters* functions were used to do unsupervised cell clustering and differential gene expression (*FindAllMarkers*) was used to find mare genes for cluster annotation. The results were then visualised using UMAP dimensionality reduction. Gene set enrichment analysis was carried out using gProfiler2 R package [46]. For Pseudo-bulk sample analysis, the seurat object was split by CAR variant metadata annotation and the *AverageExpression* Seurat function was used on the counts slot of each object, next DeSeq2 package was used to do PCA analysis and gene clustering. Gene set scores were computed using Ucell R package [47].

### Cytokine secretion ELISA

The cytokine secretion of CAR T cells was measured following co-culture with SKBR3 target cells. For each replicate, 80 000 CAR T cells and 10 000 target cells were incubated in a volume of 200 µL of culture medium for 5 days. The culture supernatant was obtained by centrifugation and used with the Bio-Plex Pro Human Cytokine 8-plex Assay (Bio-rad) according to manufacturer’s instructions. Briefly, supernatants were co-incubated with washed magnetic fluorescent beads coated with capture antibodies for the analytes GM-CSF, IFN-γ, IL-2, IL-4, IL-6, IL-8, IL-10 and TNF-α. Following washes, beads were co-incubated with PE-labeled antibodies specific to the analytes. After final washes, beads were analyzed using a Bio-Plex MAGPIX (Bio-rad). IL-2 data were excluded from the analysis due to the supplementation of this cytokine in the medium.

### Formation of MCF-7/GFP spheroids

Individual MCF-7/GFP spheroids were formed in ultra-low adherent, Nunclon™ Sphera™ U-shaped-bottom, 96-well plate (Thermo Fisher Scientific). In brief, cells were detached from the cell culture flask using 1X TrypLE™ Express enzyme (Gibco) and re-suspended at a density of 10^4^ cells/mL in pre-warmed complete minimum essential media (MEM) that contains 5 mg/mL human recombinant insulin (Gibco), 1X MEM non-essential amino acids (NEAA) (Gibco), 1 mM sodium pyruvate (Merck), and 50 mg/mL Kanamycin (BioConcept). 100 mL of cell suspension was loaded into each U-shaped-bottom well, resulting in an initial seeding of 1000 cells/spheroid. Cells were spun down at 250 g for 2 min and then kept in a humidified incubator at 37 °C and 5% CO_2_ (Binder GmbH) without medium exchange for 3 days. Before each experiment, MCF-7 spheroids were imaged with a Cell3iMager Neo plate scanning system (SCREEN Group) for quality check. Compact MCF-7/GFP spheroids with a diameter of approximately 350 mm were qualified for further experiment.

### Live imaging of cytotoxicity

CAR T cells and GFP-expressing cancer cells were co-cultured in an incubated chamber equipped with a wide field Nikon Ti2 microscope to visualize target cell death. For SKBR3 killing, cells were mixed at designated ratios in a 96-well transparent bottom plate in X-Vivo 15 without phenol red (Lonza) supplemented with FBS (Gibco) and 100 U/mL IL-2 (Peprotech). For MCF-7 killing, microtissues of 10^3^ cells were co-cultured in MT medium supplemented with 100 U/mL IL-2. Images were captured every 8 or 12 hours for 100 to 120 hours. For image analysis, a pipeline in Cell Profiler was used to extract either the total fluorescence of detected cell objects (SKBR3) or the total field fluorescence (MCF-7). The resulting values were analyzed with R and plotted with Graphpad (Prism).

### Multi-parameter flow cytometry for immunological markers

Following 5-day co-cultures of CAR T cells and SKBR3 cells at a ratio of 8:1, cells were extracted by centrifugation and prepared for flow cytometry analysis of immunological markers. First, cells were stained for viability (Zombie Aqua, Biolegend) in PBS, washed and stained in FACS buffer (2% FBS, 0.1% NaN3 in PBS) for the following markers: HLA-DR-Alexa Fluor 647 (L243), CD69-Pacific Blue (FN50), CD25-PE/Cy7 (M-A251), CD137/4-1BB-PE/Dazzle 594 (4B4-1), CD45RO-Alexa Fluor 700 (UCHL1), CCR7-APC/Cy7 (3D12), CD27-BV570 (O323), CD39-FITC (A1), CD127-PE (A019D5), CTLA-4-BV785 (L3D10), LAG-3-BV711 (11C3C65), TIGIT-BV421 (A15153G) from Biolegend; CD3-BUV395 (UCHT1 clone), CD4-BUV496 (SK3), CD8-BUV805 (SK1), CD62L-BV650 (SK11), PD-1-BB700 (EH12.1), TIM3-BV480 (7D3) from BD Biosciences; and CD45RA-APC (MEM-56) from Thermo Fisher. After washing, cells were analyzed using the Cytek Aurora full spectrum flow cytometry technology. Data were further processed with FlowJo 10 software (BD Biosciences).

## DATA AVAILABILITY

### AUTHOR CONTRIBUTIONS

R.B.D.R, R.C.R. and S.T.R. designed the study; R.B.D.R., R.C.R., F.S.S. and D.P. performed experiments; O.T.P.N. and E.K. assisted with obtaining biological material. R.B.D.R., R.C.R. and S.T.R. discussed results. R.B.D.R. and R.C.R. wrote the manuscript with input and commentaries from all authors.

## ACKNOWLEDGMENTS

We thank the ETH Zurich D-BSSE Single Cell Unit for excellent support and assistance throughout this study. This work was supported by a ETH Zurich Post-doctoral Fellowship, Switzerland (to R.B.D.R.), Helmut Horten Stiftung, Switzerland (to S.T.R.) and NCCR Molecular Systems Engineering, Switzerland (to S.T.R.).

