## Supplemental Figures and Tables for "speedingCARs: accelerating the engineering of CAR T cells by signaling domain shuffling and single-cell sequencing"

### SUPPLEMENTARY INFORMATION

SUPPLEMENTARY FIGURES 1 – 8

SUPPLEMENTARY TABLES 1 - 2

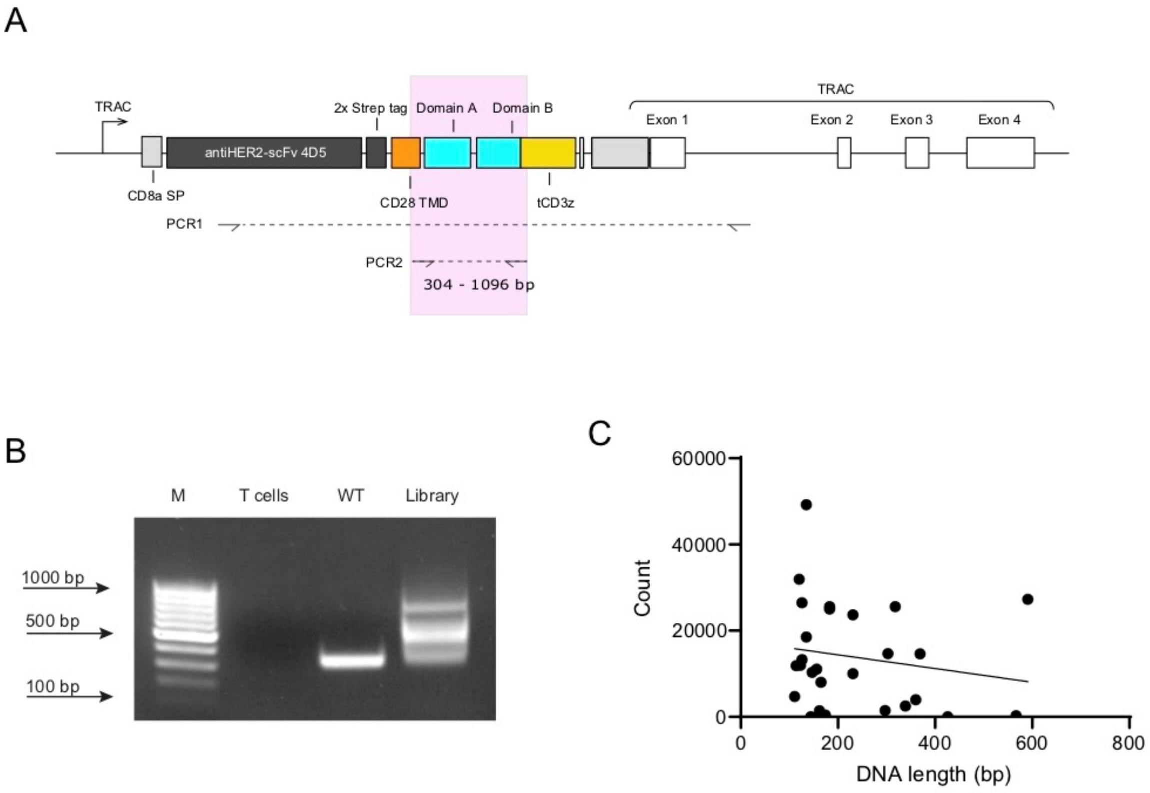


**Supplementary Figure 1: PCR amplification confirms the integration of the CAR library. A)** Schematic representation of the PCR amplification strategy to obtain amplicons of the integrated CAR gene. **B)** Agarose gel electrophoresis of the genomic amplicons from primary T cells (T cell), 28z CAR (WT) and sorted CAR T cells expressing the library of signaling domain variants (Library), all run alongside a DNA molecular weight marker (M). The library lane shows the range of expected amplicons owing to different signaling domain sizes. **C)** Scatter plot of the signaling domain's DNA length against their abundance in the CAR T cell library (post transfection, post sort). There is no significant correlation between the two variables.

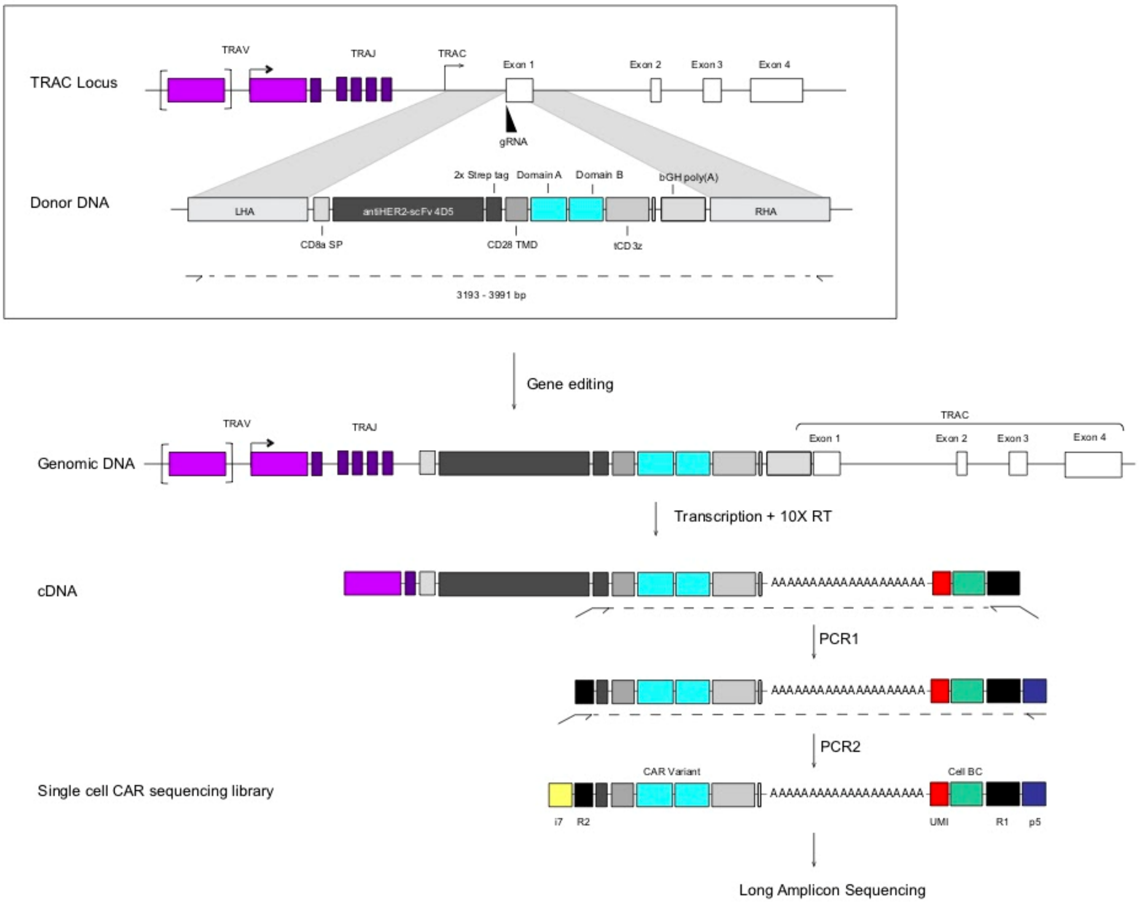


**Supplementary Figure 2: scCAR-seq PCR amplification strategy for library de-multiplexing.**

Following CRISPR-Cas9 based gene editing CAR T cells incorporate a single copy of a CAR gene into the TRAC locus. Guided by TRAC specific gene expression regulation the CAR gene is transcribed into mRNA and translated into a CAR, which is translocated to the surface of the cell to exert its immunosurveillance functions. During 10X droplet encapsulation, reverse transcription and cDNA synthesis each CAR transcript incorporates a barcode specific to its cell of origin (cell-BC). A two-step PCR amplification strategy that makes use of the synthetic Strep tag sequence found in the CAR gene, allows to selectively amplify cell-BC linked CAR transcripts from the 10X cDNA mix. This scCAR-seq library can then be sequenced using long amplicon technologies (PacBio) to trace the origin of CAR transcripts to each individual cell identified in the 10X gene expression pipeline.

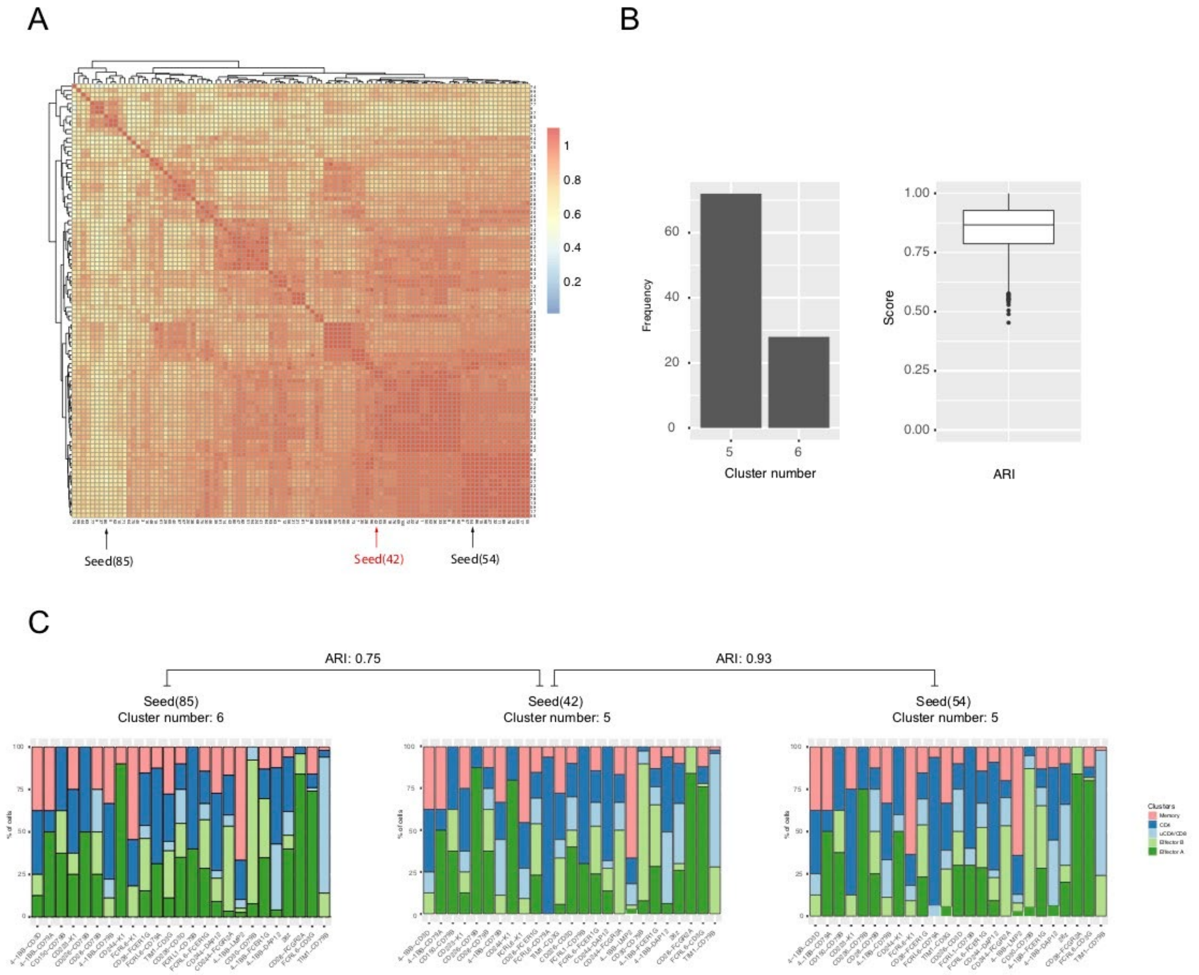


**Supplementary Figure 3: scRNA-seq clustering analysis is robust to cell subsampling. A)** Matrix of the adjusted Rand index (ARI) of the first 100 seed outputs for the clustering analysis featured in Figure 3 of the main text. This analysis relies on the random subsampling of 50 cells from CAR variants with larger cell numbers to balance the dataset. We tested how likely the subsampling can result in aberrant clustering by simulating 100 clustering procedures and measuring the similarity of the top differentially expressed genes using the ARI in a pairwise fashion. **B)** The number of clusters identified by the procedure throughout the 100 seeds is either 5 or 6. The distribution of ARIs is centered around 0.85 (lower and higher quartile 0.82 and 0.95 respectively). **C)** Examples of clustering annotation for random subsampling of the most diverging seeds according to ARI scores (seeds with highest and lowest average scores) and seed 42 used in the main analysis. Both seeds show strong similarity in the cluster's representation across the top 24 most represented CAR variants.

A

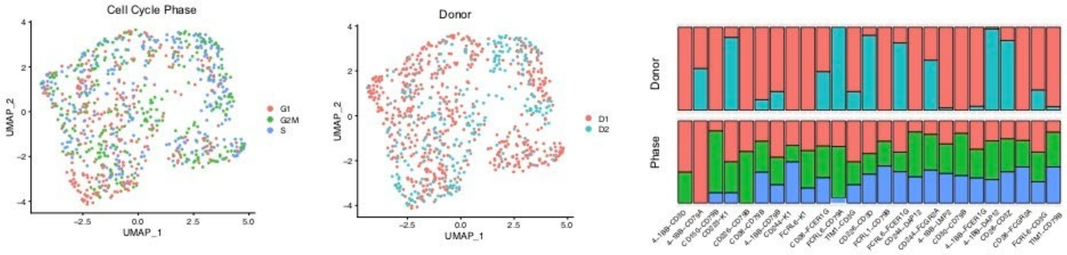


B

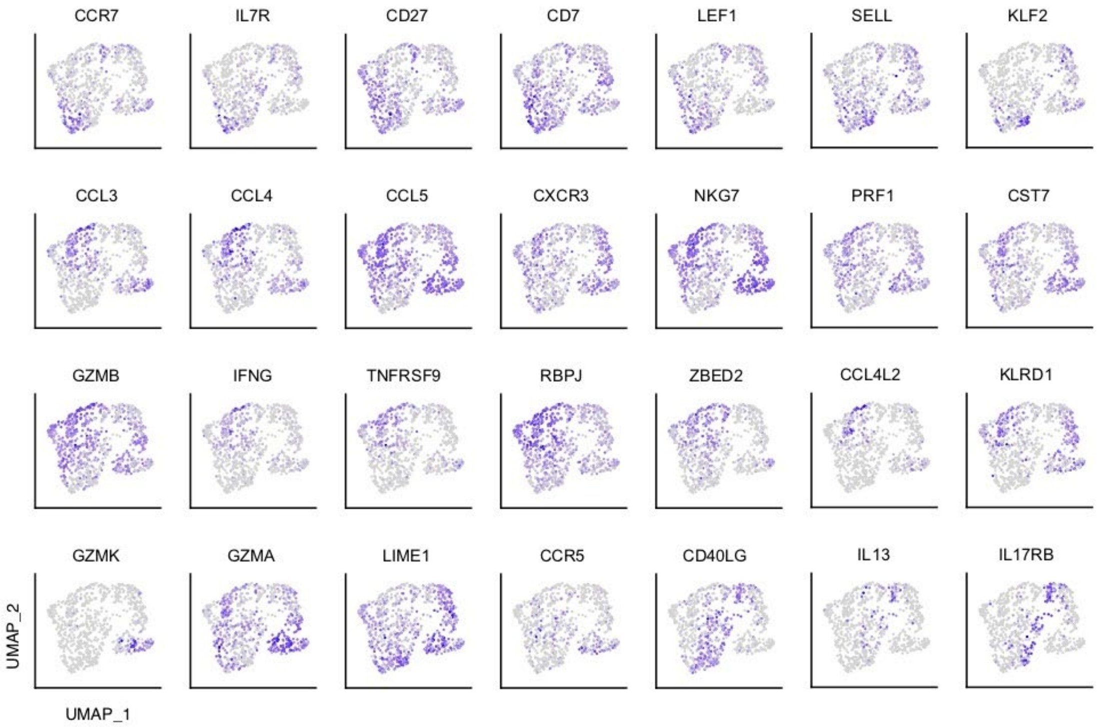


**Supplementary Figure 4: Single cell analysis of pooled library CAR T cells following tumor cell co-culture.**

**(A-C)** Pooled stimulated library CAR T cell scRNA-seq data after randomly subsampling a maximum of 50 cells per variant (seed 42). **A)** Distribution of cells based on cell donor and cell cycle phase overlaid on its UMAP embedding and as percentage of the top 25 most represented CAR variants. **B)** Feature plots showing the distribution of expression of a gene selection across the UMAP. **C)** UMAP cell annotation of the top 25 most represented CAR variants

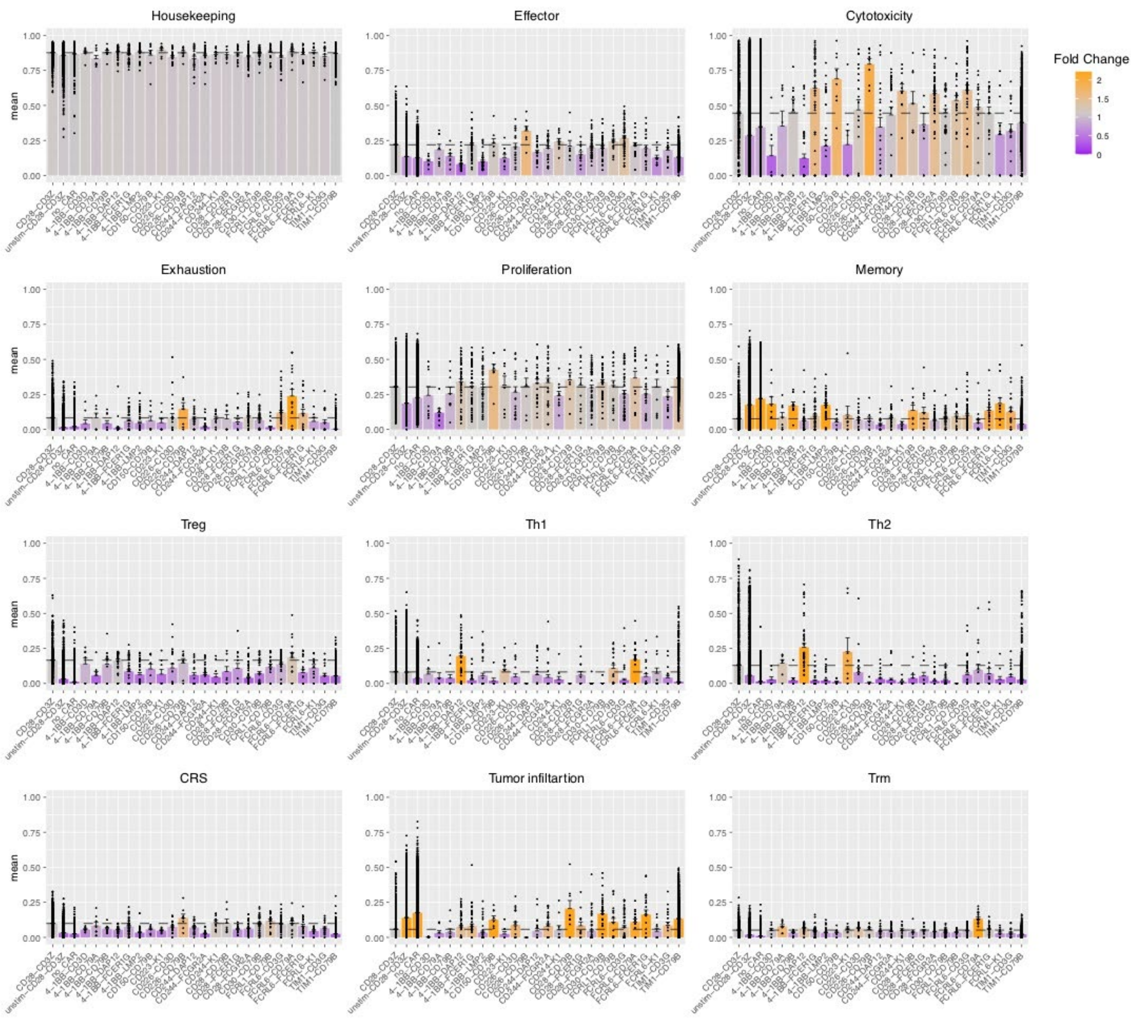


**Supplementary Figure 5: Phenotypic characterisation of CAR variants based on gene set expression scoring.**

Plots illustrating the differences in gene set scores between the 24 most represented CAR variants, stimulated and unstimulated 28z CAR and TCR- T cells, for 12 different gene sets. Dots represent the score value for each single cell (computed using Ucell), bars show the mean score across each CAR variant and colour indicates the average mean fold change of each sample group compared to stimulated 28z CAR T cells.

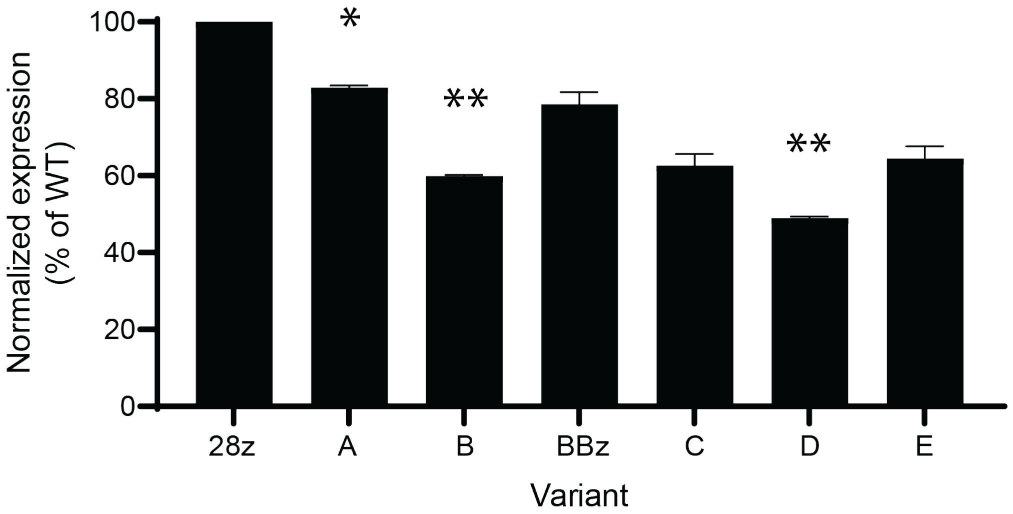


**Supplementary Figure 6: The CAR signaling variants are generally expressed at a slightly lower level on the surface of primary T cells than CAR CD28-CD3ζ ("A").** Bar plot of CAR surface levels from genome-edited primary T cells, as assessed by flow cytometry and detection of the Strep tag II. To assess significant differences between each variant and 28z, Dunnett's multiple comparisons test was used with the following significance indicators: * $P < 0.05$, ** $P < 0.001$

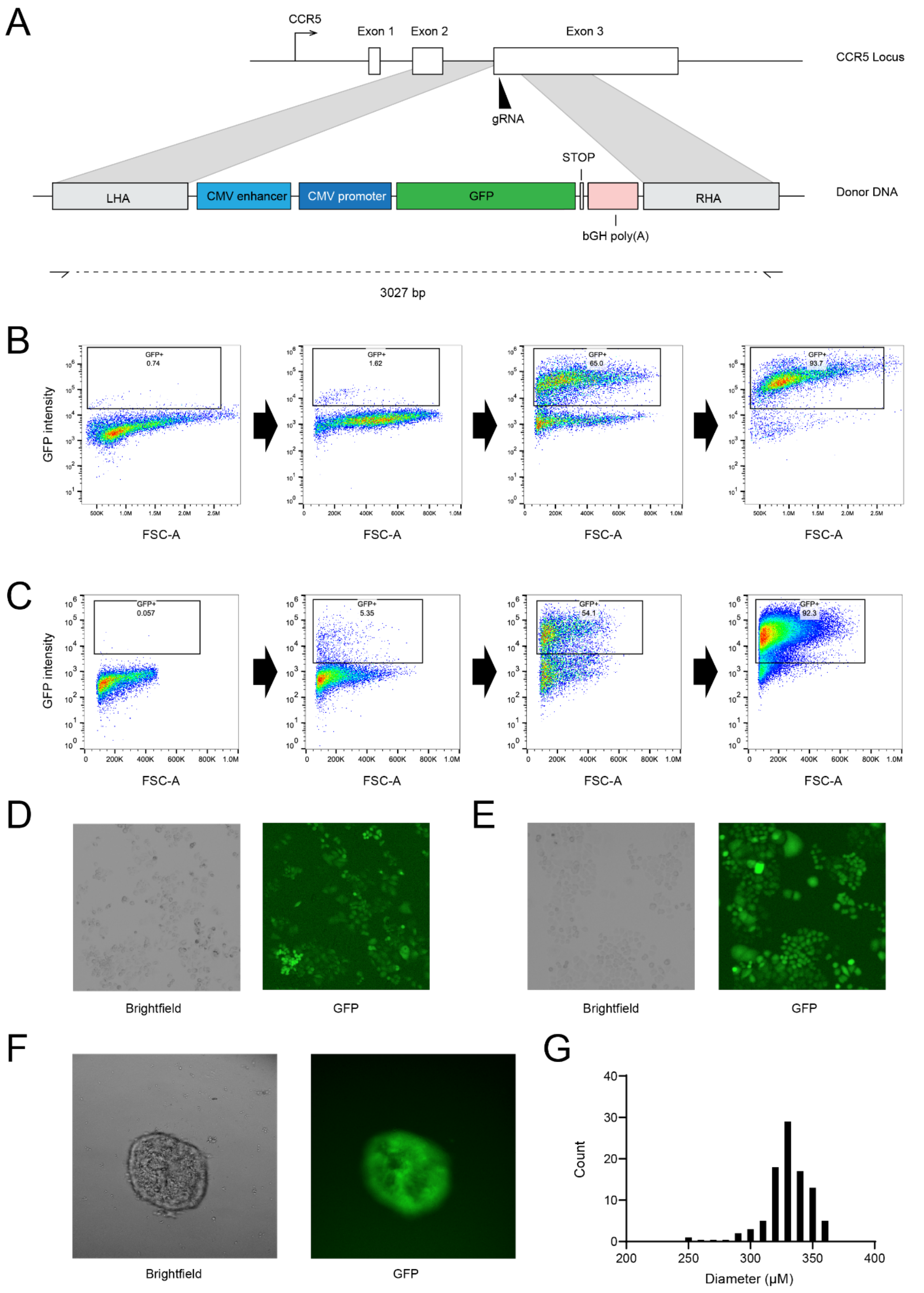


**Supplementary Figure 7: Genomic engineering of the cancer cell lines for imaging-based cytotoxicity assays. A)** The CRISPR-Cas9-based strategy for integrating a GFP expression cassette in the genome of the cell lines SKBR3 and MCF-7. A gene expression cassette was constructed harboring the two-part cytomegalovirus (CMV) promoter, the GFP ORF and a polyadenylation signal. This cassette was flanked by DNA regions homologous to the *CCR5* genomic locus and amplified by PCR to generate the repair template for HDR. A guide RNA targeting the beginning of the third exon of *CCR5* was used to generate Cas9 RNP and transfected in target cells alongside the repair template. **B) and C)** The transfected SKBR3 and MCF-7 cells respectively were sorted by FACS iteratively to select GFP-expressing cells and obtain a mostly pure (>90%) population. **D) and E)** Fluorescence microscopy confirmed the clear visibility of SKBR3 and MCF-7 cells respectively. **F)** Fluorescence image of MCF-7-GFP cells forming a spheroid structure. **G)** Frequency distribution histogram of the diameters in microns of 93 MCF-7 spheroids three days after seeding (the start of live imaging experiments).

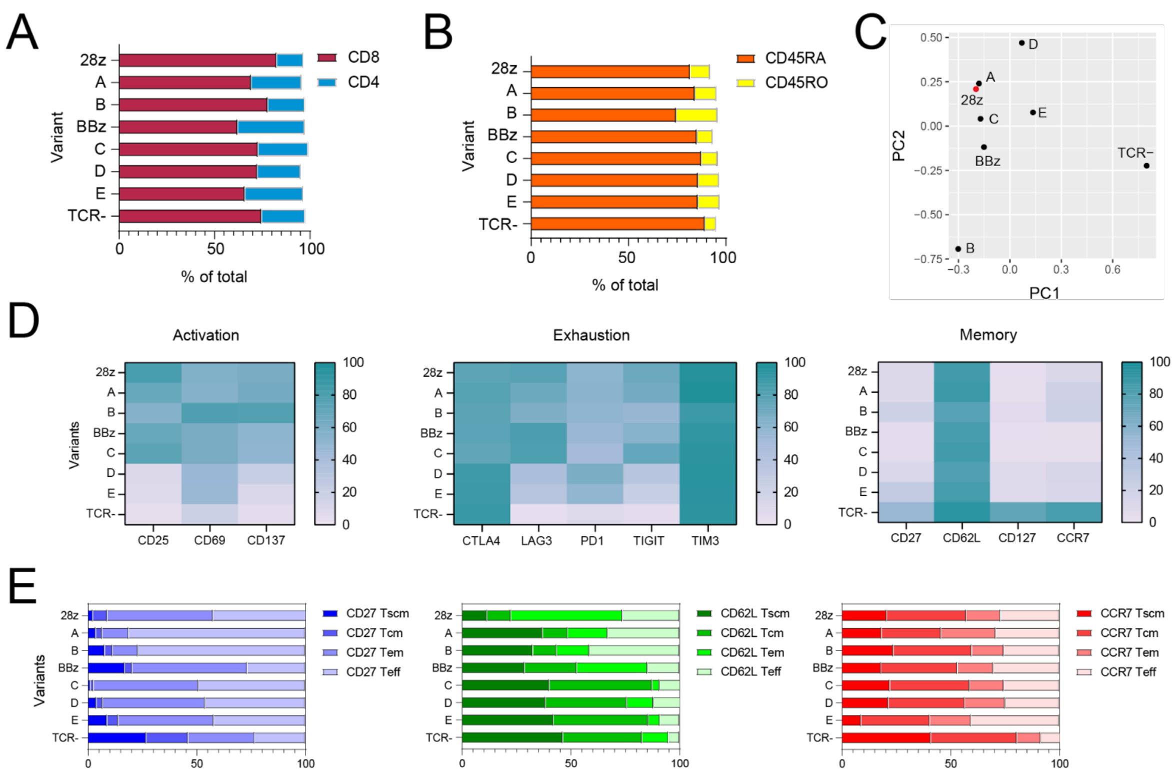


**Supplementary Figure 8: Specific subsets of CAR T cells expressing signaling domain variants based on immunological cell surface markers. A)** The proportion of CD4-positive and CD8-positive CAR T cells following co-culture is broadly similar across signaling domain variants. **B)** The proportion of CD45RO and CD45RA-positive CAR T cells following co-culture was used to subset cells into differentiation states. **C)** Scatter plot of a principal component analysis (PCA) of the surface marker expression levels of the signaling domain variants assayed. **D)** Surface expression levels of activation, exhaustion and memory cell surface markers found specifically on CD8-positive CAR T cells. The expression profiles are in broad agreement to those for all CAR T cells that are presented in Figure 5E. **E)** Comparison of alternative T cell subset definitions and the proportion of CAR T cells belonging to each subset for each variant. Subsets are defined by the presence or absence of the indicated marker: CD27, CD62L or CCR7. For all panels, surface marker expression levels were assessed by labeled-antibody staining and multi-parameter flow cytometry.

SUPPLEMENTARY TABLES

Supplementary Table 1: crRNA sequences used in this study

| Genomic target | crRNA sequence (5' to 3') |
|---|---|
| *TRAC* | CAGGGUUCUGGAUAUCUGU |
| *CCR5* | UGACAUCAAUUAUUAUACAU |

Supplementary Table 2: Oligonucleotides used in this study

| Name | Sequence (5' to 3') | Purpose |
|---|---|---|
| F1 | gttacaggCACCTGCaacaGGTG | To amplify domains from pool A for cloning |
| R1 | ggaactccCACCTGCcttgTGCTga | |
| F2 | ttagcccaCACCTGCgggcAGCA | To amplify domains from pool B for cloning |
| R2 | ccgagggcCACCTGCtcatGCGG | |
| F3 | CGGGACTAGTGGCgtcGGTTCTGGATATCTGTGGGCTGCC AGAGTTATATTGCTGGGGTT | To amplify HDR repair template with tCTS for CRISPR-Cas9 genome editing. |
| R3 | CACTTCCAGCACCgtcGGTTCTGGATATCTGTGGGCGAGA CCACCAATCAGAGGAGTTT | |
| F4 | GCTTGCTAGTAACAGTGGCCTTTAT | To amplify the recombined A and B domains within the CAR gene for sequencing |
| R4 | TACAGGCCTTCCTGAGGGTTCTT | |
| F5 | GGTCAGACAAGCTCCCGGAAAAGGA | To amplify the cytoplasmic region of the CAR gene in the *TRAC*locus for sequencing |
| R5 | AGGTGTCCCTTCCCTGCTT | |
| F5 | AATGATACGGCGACCACCGAGATCTACACTCTTTCCCTACAC GACGCTC | scCAR-seq PCR1 |
| R5 | GTGACTGGAGTTCAGACGTGTGCTCTTCCGATCTCACACCCT CAGTTCGAAAAGAGTGC | |
| F6 | AATGATACGGCGACCACCGAGATCT | scCAR-seq PCR2 |
| i7-Read2 | CAAGCAGAAGACGGCATACGAGATNNNNNNNNGTGACTGG AGTTCAGACGTGTGCTCTTCCGATC | |
